# Astrocytes mediate the dopaminergic modulation of tonic GABAergic signaling in substantia nigra

**DOI:** 10.1101/2024.03.27.586699

**Authors:** DeNard V Simmons, Oscar Andrés Moreno-Ramos, Divya D.A. Raj, Konstantin Kaganovsky, Arin Pamukcu, Nora Vrieler, Tatiana Tkatch, Zhong Xie, Ana Isabel Silva de Sousa, Enrico Zampese, Jun Ding, Rajeshwar Awatramani, Charles J. Wilson, D. James Surmeier

**Affiliations:** Department of Neuroscience, Feinberg School of Medicine, Northwestern University, Chicago, Illinois 60611, USA; Department of Neurology, Feinberg School of Medicine, Northwestern University, Chicago, Illinois 60611, USA; Department of Neurosurgery, Stanford University School of Medicine, Stanford, CA 94305; Department of Neuroscience, Developmental and Regenerative Biology, University of Texas at San Antonio, San Antonio, Texas 78249; Aligning Science Across Parkinson’s (ASAP) Collaborative Research Network, Chevy Chase, MD 20815

## Abstract

Recent studies have implicated dopaminergic signaling within the substantia nigra pars reticulata (SNr) in the emergence of Parkinsonian motor deficits. To better understand the mechanisms underlying this dependence, the ability of dopamine to modulate SNr neuron activity was studied in *ex vivo* mouse brain slices. In addition to presynaptically inhibiting phasic GABA release, D2 dopamine receptor (D2R) signaling unexpectedly suppressed a tonic, GABA_A_ receptor-mediated inhibition of SNr neuron spiking. Our studies demonstrated that this tonic modulation was largely mediated by GAT-1-dependent GABA release from ALDH1A1-expressing dopaminergic neurons. In contrast, the ability of D2R agonists to disinhibit SNr neurons depended upon D2R-expressing astrocytes and stimulation of GAT-3 uptake of GABA from the extracellular space. In addition to underscoring the importance of dendritically released dopamine in modulating synaptic transmission, our studies demonstrate that dopaminergic modulation of astrocytes plays a key role in modulating SNr circuits and motor behavior.

**Teaser:** SNc dopaminergic neurons and astrocytes regulate tonic GABAergic inhibition of SNr neurons.

## Introduction

Parkinson’s disease (PD) is the second most common neurodegenerative disease (*1, 2*). The cardinal motor symptoms of the disease – bradykinesia and rigidity – are attributable to the degeneration of dopaminergic neurons in the substantia nigra pars compacta (SNc) that innervate the basal ganglia. Although the classical model of the network dysfunction underlying PD motor symptoms emphasizes the importance of the dopaminergic innervation of the striatum (the largest of the basal ganglia nuclei), recent work has demonstrated that other basal ganglia nuclei also are important in the transition to the parkinsonian state, including the substantia nigra pars reticulata (SNr) (*3, 4*).

From the network standpoint, the SNr is particularly interesting. The principal SNr neurons are autonomously active GABAergic neurons that tonically inhibit motor control circuits in the brainstem and diencephalon (*5*). The basal activity of SNr neurons is bidirectionally regulated by synaptic connections with other basal ganglia neurons, including GABAergic neurons in the GPe and striatum (direct pathway spiny projection neurons) and glutamatergic neurons in the subthalamic nucleus (STN) (*6, 7*). These synaptic connections are generally thought to be modulated by dopamine through presynaptic G-protein coupled receptors (GPCRs). The dopamine that acts on these receptors is released locally by the dendrites of SNc neurons that course through the SNr (*8, 9*). Previous studies have suggested that dopamine released into the SNr suppresses synaptic transmission arising from GPe and STN neurons, while enhancing transmission from direct pathway spiny projection neurons (dSPNs). However, the neuropil in this region is complex, making a definitive assessment of how dopamine modulates these pathways problematic with conventional electrical stimulation methodologies.

Optogenetic methods offer a way of surmounting this difficulty, but this approach has not been used to determine how dopamine modulates either GPe or STN input to SNr neurons. Sorting this out in a rigorous way is critical to understanding not only how dopamine modulates SNr activity in the healthy brain, but how this activity goes awry in PD. The potential impact of this local dopaminergic modulation has been driven home by the observation that in parkinsonian MCI-Park mice, restoration of dopaminergic signaling within the SNr alone alleviates open field deficits (*10*).

Recent work points to another way in which dopaminergic neurons in the SNc might modulate SNr activity. Optogenetic stimulation of genetically defined dopaminergic neurons has revealed that some of these neurons co-release glutamate and others co-release GABA from their axon terminals (*11–13*). It is unclear whether release of these transmitters is a phenomenon limited to axons or whether it extends to the dendrites of SNc dopaminergic neurons.

The goal of our study was to define how dopamine modulates the activity of SNr neurons. The initial experiments utilized optogenetic approaches to characterize how dopamine modulated the release of GABA from GPe synaptic terminals on SNr neurons. These studies were consistent with inferences drawn from previous work that presynaptic D2 dopamine receptors (D2Rs) inhibit phasic release of GABA from GPe terminals. However, these experiments also revealed that D2Rs suppressed a tonic GABAergic inhibition of SNr spiking. This tonic GABAergic signal was traced back to a subset of SNc dopaminergic neurons that express aldehyde dehydrogenase 1α1 (ALDH1A1). The dendritic release of GABA was dependent upon GAT-1 transporters and glucose availability, suggesting it reflected metabolic state, not spiking *per se.* In contrast, the suppression of tonic GABA currents was largely dependent upon a subset of D2R-expressing SNr astrocytes and stimulation of GABA uptake by GAT-3 transporters. The modulation of SNr astrocytic GABA uptake establishes an unexpected way in which dendritic dopamine release modulates local circuitry, potentially increasing the dynamic range of SNr signaling to downstream motor circuits.

## Results

### D2R agonists inhibited phasic GABA release from GPe terminals

To characterize how dopamine modulated GABA release from GPe terminals, GPe neurons were induced to express the opsin Chronos by injecting an adeno-associated virus (AAV) carrying a synapsin promoter driven expression plasmid with a tdTomato fluorescent protein reporter (see Methods) into the GPe of mice (Fig. 1A). Optogenetic stimulation of GPe axons evoked robust post-synaptic currents in SNr neurons voltage clamped at -60 mV with a Cs^+^ containing whole-cell electrode (Fig. 1B). These optically evoked currents were blocked by bath application of the GABA_A_ receptor (GABA_A_R) antagonist gabazine (10 µM). To assess dopaminergic modulation of synaptic transmission, the D2R agonist quinpirole (2 µM) was bath applied while optogenetically stimulating GPe axons. Quinpirole reduced the amplitude of the GPe evoked inhibitory postsynaptic currents (IPSCs) by roughly 30% (p = 0.02, n = 5) (Fig. 1 B, C). Consistent with a presynaptic locus of the modulation, the paired-pulse ratio of GPe evoked IPSCs rose significantly in the presence of quinpirole (p = 0.02, n = 6) (Fig. 1 D, E).

**Figure 1.**
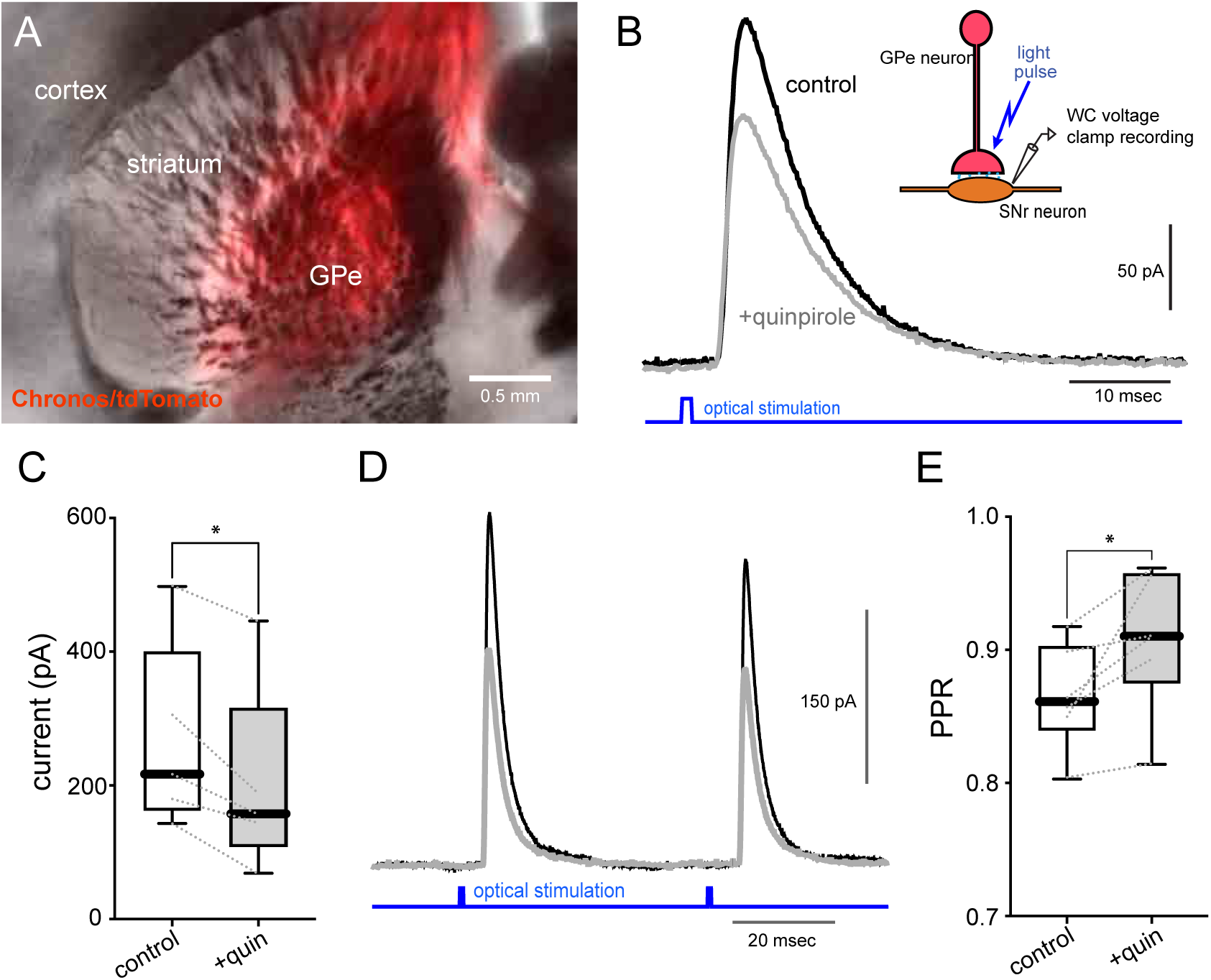
Optically Evoked GPe mediated IPSCs are depressed by D2 Receptor activation. (A) Coronal slice, injection site of AAV-Chronos/tdTomato into GPe. (B) Photo-evoked GPe mediated IPSCs recorded in whole cell configuration from SNr neurons. (C) Paired T-test of photo-evoked IPSC amplitudes before and after bath application of 2 μM quinpirole; n = 5, p = 0.02. (D) Paired pulses of GPe-mediated IPSCs at 50 ms interval before and after quinpirole bath application. (E) Paired T-test of paired pulse ratio of GPe-mediated IPSCs before and after quinpirole application; n = 6, p = 0.02

To assess the impact of this presynaptic modulation on the ability of GPe to modulate the basal spiking of SNr neurons, SNr neurons were recording in cell-attached mode (to preserve the intracellular environment) while GPe axons were stimulated at 30 Hz for 2 seconds. Initially, this stimulation effectively suppressed SNr activity, but the efficacy of this inhibition waned with time (Fig. 2A-C). The waning strength of the GPe inhibition is likely attributable to both biologically relevant factors, like depression of GABA release, but also to the limitations posed by the optogenetic methods (*14*). As expected, quinpirole attenuated the ability of GPe stimulation to slow the spiking of SNr neurons (p = 0.02, n = 8) (Fig. 2A-C). Taken together, these results are consistent with the inferences drawn from previous work using electrical stimulation (*15*).

**Figure 2.**
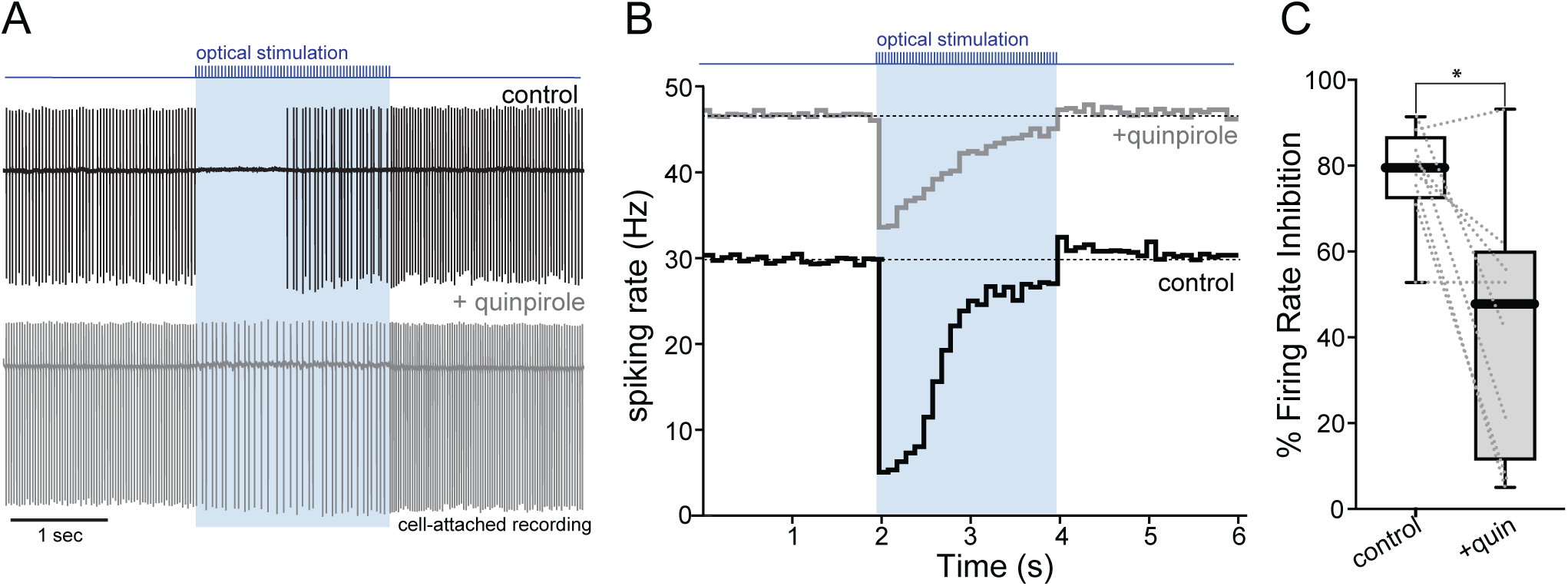
Train stimulation of GPe input inhibits SNr spiking. (A) A 6 second cell attached recording of SNr spiking inhibited by a 2 sec 30 Hz train stimulation of GPe-mediated IPSCs, before and after bath application by quinpirole. (B) A 6 second PSTH of cell attached spiking inhibited by a 2 sec 30 Hz train stimulation of GPe-mediated IPSCs, before and after bath application of quinpirole. (C) Wilcoxon matched-pairs signed rank test of percent inhibition (change in discharge rate / basal discharge rate) before and after quinpirole, n = 8, p = 0.02

### D2R agonists lowered tonic GABA release

One of the unexpected observations in these experiments was the elevation in the basal spiking rate of SNr neurons with bath application of quinpirole, in the absence of GPe axon stimulation (p < 0.01, n = 8) (Fig. 3A). The quinpirole-induced elevation in discharge rate was about 50% when monitored in cell-attached mode. Bath application of gabazine (10 µM) increased the basal spiking rate of SNr neurons recorded in cell-attached mode by roughly 25%, suggesting that they were tonically inhibited by activation of GABA_A_Rs (p < 0.01, n = 10) (Fig. 3B). These results suggest that quinpirole and gabazine might be acting through a common mechanism – inhibition of tonic GABA release. If this is the case, bath application of gabazine should occlude the modulation of SNr spiking by quinpirole. Indeed, in the presence of gabazine, quinpirole did not significantly modulate SNr spiking rate (p = 0.13, n = 7) (Fig. 3C).

**Figure 3.**
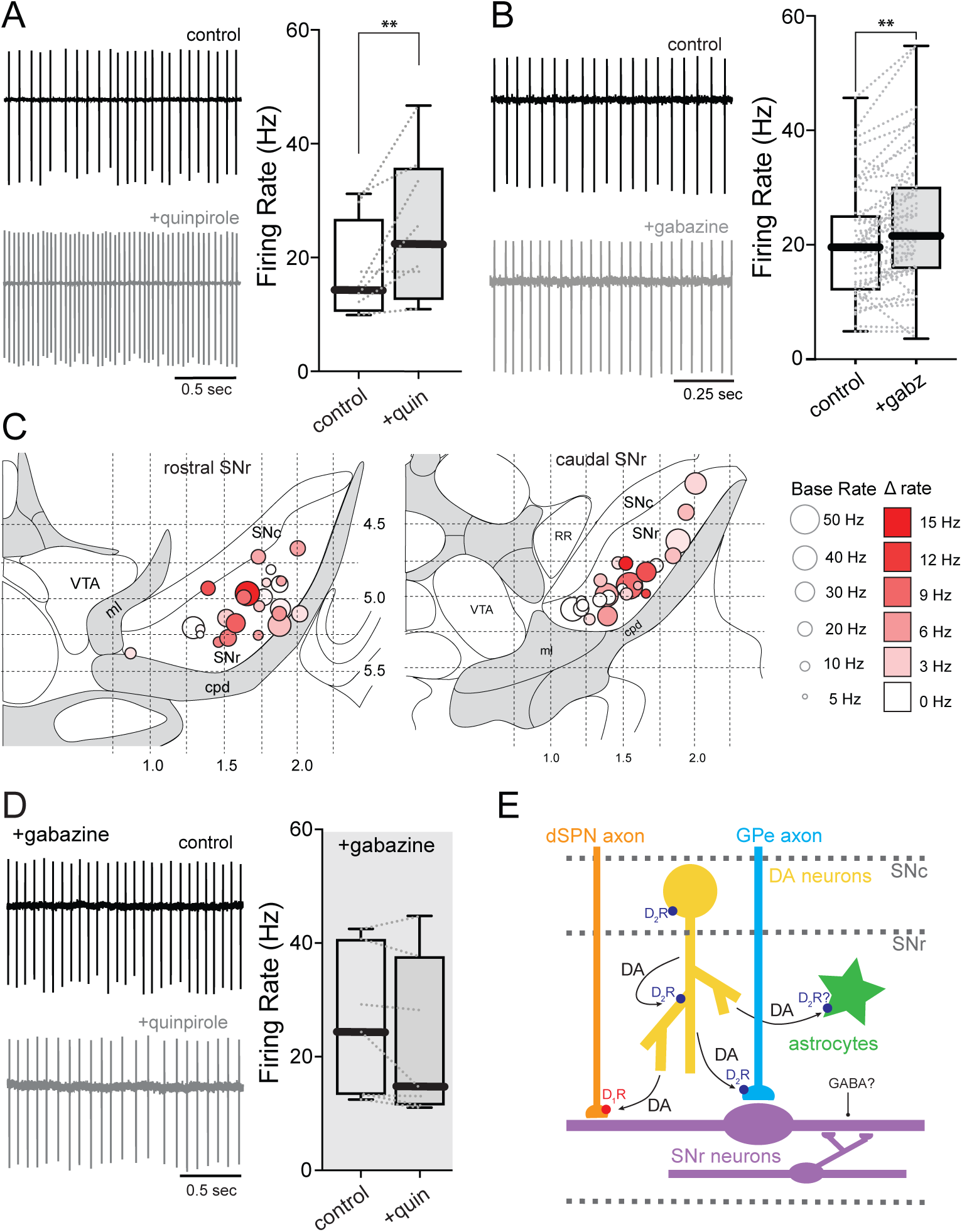
GABA-A receptor blockade occludes disinhibition by quinpirole. (A) Example trace and summary data of cell attached spiking from SNr neurons before and after bath application of quinpirole. Paired T-test, n = 8, p < 0.01. (B) Example trace and summary data of cell attached spiking from SNr neurons before and bath application of gabazine. Paired T-test, n = 45, p < 0.01. (C) Rostral and Caudal maps of gabazine treated cells recorded in SNr. Circle diameter represents initial firing rate. Circle hue represents absolute change in rate post gabazine bath application. (D) Example trace and summary data of cell attached spiking from SNr neurons pretreated with gabazine, before and after bath application of quinpirole. Paired T-test, n = 7, p = 0.06. (E) Hypothesized schematic of cells in SNr and SNc, showing the expression of D1 and D2 receptors that could be modulating GABA release upon SNr neurons.

These results demonstrate that there is a tonic source of GABA that significantly modulates the spiking rate of SNr neurons and is negatively modulated by D2-class receptors. Within the SNr, there are several possible sources of tonic GABA release that might be shaped by D2R signaling (Fig. 3E): GPe terminals, astrocytes, dSPN terminals and SNc dopaminergic neurons. As SNr neurons are not known to express D2-class dopamine receptors, they were excluded as possible sources of tonic GABAergic signaling.

### Tonic GABA release was not derived from GPe terminals or astrocytes

One potential source of tonic GABA release that might be negatively modulated by D2Rs is the GPe terminal. To test this hypothesis, an AAV carrying a synapsin-driven hM4D (Gi) designer receptor exclusively activated by designer drugs (Gi-DREADD)-tdTomato expression construct was injected into the GPe. Bath application of the DREADD agonist CNO reduced the amplitude of electrically evoked IPSCs in voltage-clamped SNr neurons (p = 0.02, n = 4) (Fig. 4A). This modulation was similar in magnitude to that seen in optogenetically evoked IPSCs following D2R stimulation.

**Figure 4.**
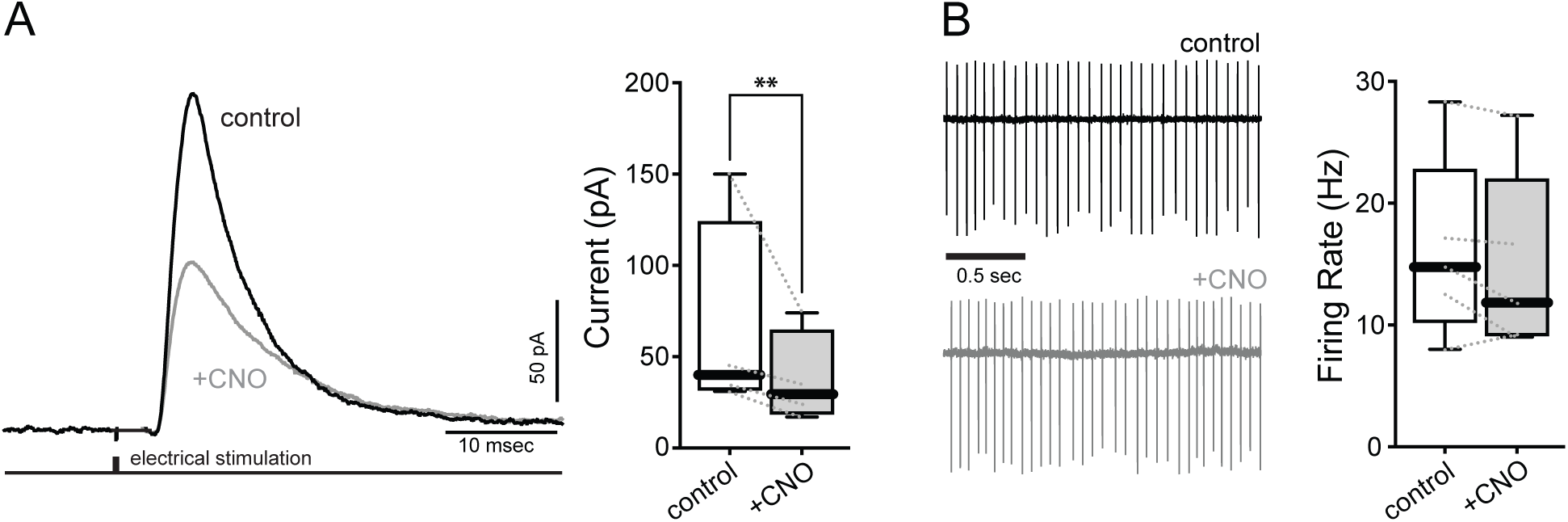
Inhibition of GPe input does not alter tonic spiking in SNr. GPe injected with inhibitory Gi-DREADDs. (A) Example trace and summary data of electrically evoked GPe-mediated IPSCs before and after bath application of DREADD agonist CNO. Paired T-test, n = 4, p < 0.01. (B) Example trace and summary data of cell attached spiking from SNr neurons before and after bath application of CNO. Paired T-test, n = 5, p = 0.16

However, CNO application had no effect on the basal spiking rate of SNr neurons recorded in cell-attached mode, arguing that GPe terminals were not contributing to tonic GABA release (p = 0.19, n = 5) (Fig. 4B).

In other regions of the brain, astrocytes express D2Rs (*16–18*). Moreover, in some regions of the brain, astrocytes can release GABA (*19*). Astrocytic GABA release is thought to occur through bestrophin1 channels in response to an elevation in intracellular Ca^2+^ concentration (*20*). To test if astrocytes contributed to tonic GABAergic signaling, an AAV carrying a plasmid in which a glial fibrillary acidic protein (GFAP) promoter fragment-controlled expression of an excitatory hM3 (Gq) DREADD construct was stereotaxically injected into the SNr of mice. These vectors have been used to chemogenetically elevate intracellular Ca^2+^ in astrocytes (*21*) (Fig. S4A). In these slices, bath application of the DREADD agonist CNO elevated the basal spiking rate of SNr neurons recorded in cell-attached mode (p = 0.03, n = 8) (Fig. S4B). In line with the notion that astrocytes were exciting (not inhibiting) SNr neurons, bath application of the bestrophin1 inhibitor NPPB decreased SNr spiking rate (p = 0.03, n = 6) (Fig. S4C).

### Dopamine enhanced tonic GABA release from dSPN terminals

In addition to suppressing GABA release from GPe terminals, dendritically released dopamine can facilitate GABA release from dSPN terminals invested with D1Rs (*22–24*). Although previous work has focused on spike-evoked GABA release, it is also possible that D1R stimulation promotes tonic GABA release from these terminals. Thus, by suppressing dendritic dopamine release, D2R stimulation might dis-facilitate GABA release from dSPN terminals. To test this possibility, a D1R antagonist was bath applied while monitoring SNr spike rate. Bath application of SCH39166 (2 µM) did indeed significantly elevate SNr spiking rate (p = 0.02, n = 12) (Fig. S2), but much less than that produced by the D2R agonist – arguing that there was another source of tonic GABA release in the SNr.

### SNc dopaminergic neurons directly contributed to tonic GABA release

Another possible source of tonic GABA release are SNc dopaminergic neurons. As mentioned above, the axons of SNc dopaminergic neurons co-release dopamine and GABA, raising the possibility that co-release also occurs from their dendrites. Although somewhat controversial, the release of GABA from the axons of SNc dopaminergic neurons has been reported to be sensitive to knockdown of ALDH1A1 (*25*). To assess the extent to which the dendrites of SNc dopaminergic neurons coursing through the SNr express ALDH1A1, mice expressing Cre recombinase in ALDH1A1+ (Aldh1a1-2A-iCre) neurons were crossed with mice expressing the recombinase FlpO under the control of the dopamine transporter promoter (DAT-2A-Flpo). These mice were then injected with AAV8 HSyn-Con/Fon-YFP, and AAV8 Ef1a-Coff/Fon-mCherry viruses. In tissue from these mice, ALDH1A1+ dopaminergic neurons were marked by EYFP expression, whereas ALDH1A1- dopaminergic neurons were marked with mCherry. In both rostral and caudal regions, the dendrites of ALDH1A1+ dopaminergic neurons coursed through much of dorsoventral extent of SNr (Fig. 5A, B) (*9, 26*).

**Figure 5.**
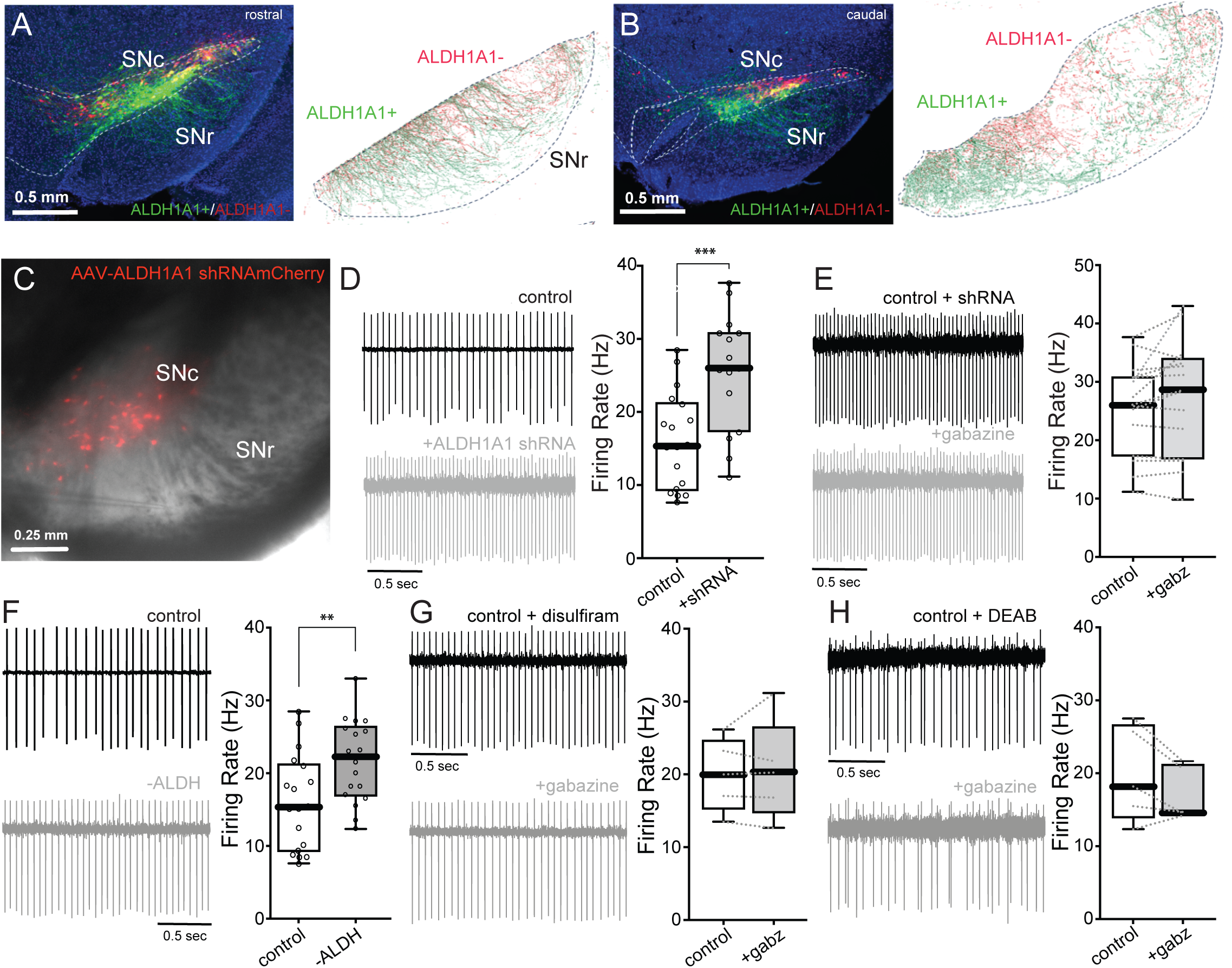
Blockade of ALDH disrupts tonic GABA in SNr. (A) Coronal image of DAT-Cre+ / ALDH1A1+ (green) and DAT-Cre+ / ALDH1A1- (red) neurons in rostral substantia nigra. (B) Coronal image of DAT-Cre+ / ALDH1A1+ (green) and DAT-Cre+ / ALDH1A1- (red) neurons in caudal substantia nigra. (C) Coronal image of ALDHA1-shRNA-mCherry expression in SNc. (D) Example trace and summary data of cell attached spiking from control SNr neurons, and SNr neurons from animals injected with ALDH-shRNA. Welch’s T-test, n = 18 control/15 shRNA, p < 0.001. (E) Example trace and summary data of cell attached SNr neuron spiking from mice injected with ALDH-shRNA before and after bath application with gabazine. Paired T-test, n = 15, p = 0.05. (F) Example trace and summary data of SNr spiking from control neurons and neurons pretreated with ALDH1A1 antagonists DEAB or disulfiram. Welch’s T-test, n = 18 control/18 blocked, p < 0.01. (G) Example trace and summary data of SNr spiking pretreated with disulfiram, before and after bath application of gabazine. Paired T-test, n = 5, p = 0.40. (H) Example trace and summary data of SNr spiking pretreated with DEAB, before and after bath application of gabazine. Paired T-test, n = 5, p = 0.10.

To determine if ALDH1A1 in these dendrites contributed to the generation of tonic GABA signaling, a viral knockdown strategy was used. The SNc of mice were stereotaxically injected with an AAV carrying an ALDH1A1 shRNA expression construct with an mCherry reporter (Fig. 5C) (*25*). In ex vivo, cell-attached recordings from SNr neurons, the basal spiking rate was significantly higher than in controls (p < 0.001, n = 18 control / 15 shRNA) (Fig. 5D). Moreover, bath application of gabazine did not significantly change the basal spiking rate of SNr neurons from ALDH1A1 shRNA treated mice (p = 0.05, n = 15) (Fig. 5E).

To provide an orthogonal test of the hypothesis that ALDH1A1 (or a closely related aldehyde dehydrogenase) was driving GABA synthesis, a pharmacological approach was used. First, the aldehyde dehydrogenase inhibitor disulfiram (10 µM)(*27*) was bath applied to control slices and then the effect of gabazine on basal spiking rate monitored.

Disulfiram blocked the effect of gabazine (p = 0.40, n = 5) (Fig. 5G). Second, another aldehyde dehydrogenase inhibitor N,N-diethylaminobenzaldehyde (DEAB, 10 µM)(*28*) was bath applied, and the effect of gabazine on basal spiking monitored. As with disulfiram, gabazine had no effect on DEAB-exposed SNr neurons (p = 0.10, n = 5) (Fig. 5H).

These experiments suggest that ALDH1A1 is involved in the intracellular generation of GABA in dopaminergic neurons. Another enzyme implicated in GABA synthesis is monoamine oxidase (MAO) (*29*). Dopaminergic neurons express both MAO-A and MAO-B (*30*). Indeed, incubating brain slices in the irreversible MAO-B antagonist rasagiline (10 µM) blunted any discernible effect of gabazine on the spiking rate of SNr neurons (p = 0.23, n = 9) (Fig. 6A). These results make the case that the tonic GABA signal in the SNr largely originated in the cytosol of dopaminergic neuron dendrites.

**Figure 6.**
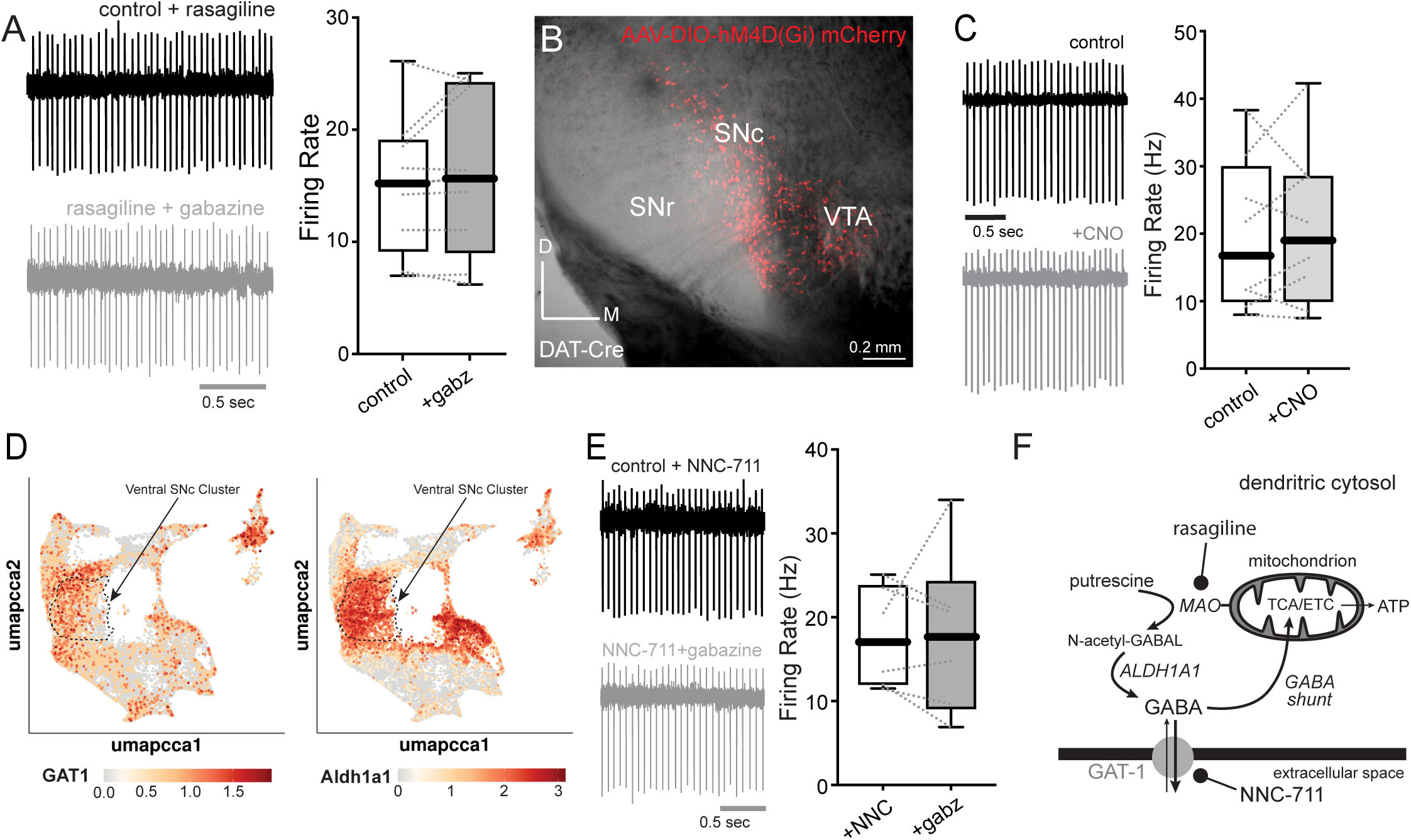
Synthesis of GABA in dopaminergic neurons is mediated by ALDH1A1, and release is mediated by GAT-1. (A) Example trace and summary data of SNr spiking pretreated with rasagiline, before and after bath application of gabazine. Paired T-test, n = 9, p = 0.23. (B) Injection site of AAV-DIO-hM4D(GI)-mCherry into the SNc of DAT-Cre mice. (C) There was no significant change in SNr neuron firing rate when CNO was bath applied (p = 0.29, n = 8). (D) GAT-1 expression is enriched in ALDH1A1+ dopaminergic neurons. (E) Example trace and summary data of SNr spiking pretreated with NNC-711, before and after bath application of gabazine. Paired T-test, p = 0.32, n = 6. (F) Schematic of non-canonical synthesis of GABA feeding into the GABA shunt at SNc dopamine neuron mitochondria.

How then might GABA have moved from the cytosol to the extracellular space? One possibility is activity-dependent vesicular release (*31, 32*). If this was the mechanism of release, then chemogenetically inhibiting them – slowing or stopping their autonomous pacemaking – should decrease GABA release, accelerating SNr spiking. To test this hypothesis, the SNc of DAT-Cre mice was injected with an AAV carrying a Cre-dependent expression construct for hM4-DREADD (Fig. 6B). Stimulation of this Gi-coupled DREADD should open Kir3 K^+^ channels in SNc dopaminergic neurons, slowing their spiking rate. Indeed, in brain slices from mice injected 3 weeks prior, bath application of the DREADD activator clozapine-N-oxide (CNO) significantly slowed the spiking rate of SNc dopaminergic neurons (mean reduction of 19%, p = 0.02, n = 6) (Fig. S3). However, CNO application did not increase the spiking rate in SNr (p = 0.29, n = 8) (Fig. 6C).

Another potential path for GABA is through the GAT-1 transporter (*33–35*). Interestingly, queries of Dopabase (*36*) revealed that GAT-1 expression was enriched in ALDH1A1+ dopaminergic neurons (Fig. 6D); roughly one-third of the ventral SNc dopaminergic neurons in clusters 4 and 10 had detectable levels of both transcripts. To determine whether GABA was being released through GAT-1, brain slices were incubated in GAT-1 antagonist NNC-711 prior to and during recording. In the presence of NNC-711, bath application of gabazine failed to increase SNr firing rate (p = 0.32, n = 6) (Fig. 6E). Taken together, these experiments support the hypothesis that GABA release from dopaminergic dendrites was mediated by GAT-1.

If GABA is being extruded through GAT-1, what biological purpose is cytosolic GABA serving in ALDH1A1+ dopaminergic neurons? ALDH1A1+ dopaminergic neurons are the most bioenergetically challenged subtype of dopaminergic neuron, manifesting high levels of mitochondrial oxidant stress (*37–39*). It would make sense if they had an alternative energy source available when glucose was in short supply.

Cytosolic GABA could be this source, acting as a substrate for the "GABA shunt", which generates succinate that can be used by the mitochondrial electron transport chain (ETC) to drive oxidative phosphorylation (OXPHOS). Indeed, ALDH1A1+ dopaminergic neurons express GABA transaminase (GABA-T) and succinic semialdehyde dehydrogenase (SSADH) – two enzymes necessary to metabolize GABA to succinate(*31, 40*). If this were the case, lowering extracellular glucose concentration should increase cytosolic GABA metabolism and shift the GAT-1 reversal potential to more depolarized membrane potentials, diminishing or eliminating GABA release (*41*). To test this hypothesis, the extracellular glucose concentration was lowered to 2.5 mM, and the sensitivity of SNr spiking to gabazine was re-examined. As predicted, in this condition, gabazine did not significantly increase SNr spiking (Fig. S8).

To better define the mechanisms underlying this shift, the contribution of GABA metabolism to OXPHOS in SNc dopaminergic neurons was examined by using a genetically-encoded fluorescent ratiometric ATP/ADP probe, PercevalHR (*42*). Two-photon light scanning microscopy (2PLSM) was used to monitor PercevalHR fluorescence ratio in dopaminergic neurons in *ex vivo* midbrain slices (*43*) (Fig.S9A). Bath application of the GABA-T inhibitor Vigabatrin (*44*) decreased the PercevalHR ratio, consistent with a drop in mitochondrial ATP generation (Fig.S9B,C). To dissect the impact of GABA-T inhibition on mitochondrial OXPHOS and cytosolic glycolysis, PercevalHR fluorescence was monitored after bath application of the complex V inhibitor oligomycin (10 µM) and then glucose replacement with the non-hydrolyzable analogue 2-deoxyglucose (2-DG, 3.5mM) (*43*). Consistent with engagement of the GABA shunt, GABA-T inhibition selectively decreased the mitochondrial contribution to maintenance of cytosolic ATP levels. These features of GABA metabolism in dopaminergic neurons are summarized in Figure 6F.

### Astrocytic D2Rs promote GABA uptake through GAT-3 transporters

If dopaminergic neurons were the source of the tonic GABA in the SNr, what were the mechanisms responsible for its D2R-mediated suppression? In other brain regions (*16–18, 45*), astrocytes express D2Rs that stimulate GAT-3 activity (*35, 46, 47*). As a first step toward determining if the same was true in the SNr, RNAScope was used to determine whether D2Rs co-localized with the astrocytic marker S100B in the SNr. Indeed, the two were co-localized in a significant fraction of both lateral and medial SNr astrocytes (lateral SNr: mean ratio of D2R positive astrocytes = 37 ± 4%; medial SNr: mean ratio of D2R positive astrocytes = 32 ± 4%; n = 3 mice/516 astrocytes) [Fig. 7A, Fig. S7]. To determine whether G_i/o_-coupled D2Rs could stimulate astrocytic GABA uptake, the SNr was stereotaxically injected with an AAV carrying a plasmid in which the expression of hM4Di was controlled by the GFAP promoter (Fig. 7B). SNr neurons were monitored in cell-attached mode. Bath application of CNO to activate hM4Di significantly increased the spiking rate of SNr neurons (28% mean increase, p < 0.01, n = 10) (Fig. 7C,D). The CNO effect on SNr spiking rate was occluded by application of gabazine – directly implicating astrocytic regulation of GABA (p = 0.10, n = 11) (Fig. 7E).

**Figure 7.**
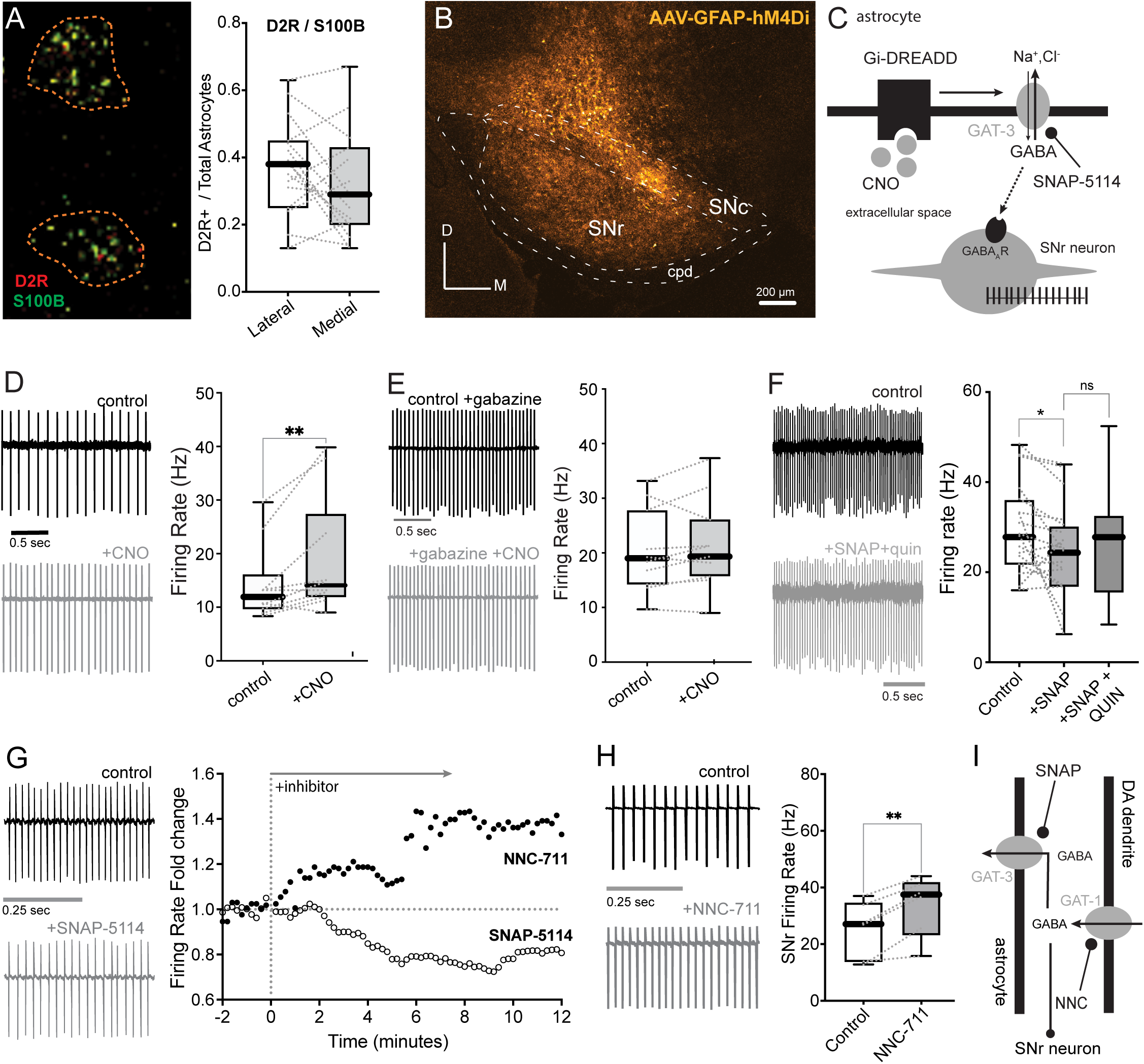
Astrocytic D2Rs promote GABA uptake through GAT-3 transporters. (A) RNAscope image of astrocytes labelled with S100B coexpressing D2Rs and summary data of D2R colocalization in SNr astrocytes. Lateral SNr: mean ratio of D2R positive astrocytes = 37 ± 4%; Medial SNr: mean ratio of D2R positive astrocytes = 32 ± 4%; n = 3 mice/516 astrocytes. (B) Injection site of AAV-GFAP-hM4D(Gi)-mCherry into the SNr of wildtype mice. (C) Schematic of Gi-DREADD activation enhancing reuptake of GABA by GAT3 in astrocytes, indirectly disinhibiting SNr neurons. (D) SNr neuron activity was recorded in cell-attached mode, and it was found that activation of Gi-DREADD in astrocytes by CNO bath application significantly facilitated SNr firing (28% mean increase, p < 0.01, n = 10). (E) There is no increase in firing rate of SNr neurons observed when gabazine was bath applied prior to CNO (p = 0.10, n = 11). (F)An example trace and summary data of SNr firing before and after bath application of SNAP-5114 and quinpirole. Bath application significantly reduces SNr firing from baseline (Paired T-test, p = 0.02, n = 19). Bath application of quinpirole after SNAP-5114 fails to significantly increase SNr firing (Welch’s T-test, p = 0.27). (G) An example trace of SNr firing before and after bath application of SNAP-5114; Example time series of SNr neurons following bath application of either NNC-711 or SNAP-5114. (H) An example trace and summary data of SNr firing before and after bath application of NNC-711 (p < 0.01, n = 6). (I) Schematic of efflux of tonic GABA from DA neuron dendrites and uptake of tonic GABA at SNr astrocytes, with the remaining extracellular GABA inhibiting SNr firing.

To provide an additional assay for astrocytic involvement, the GAT-3 selective antagonist SNAP-5114 (20 µM) (*48*) was applied to brain slices while monitoring SNr spiking rate (Fig. 7F). The GAT-3 selective antagonist significantly decreased SNr firing rate (23% decrease, p = 0.02, n = 19). In the presence of the GAT-3 antagonist, bath application of quinpirole had no effect on SNr spiking rate (p = 0.27) (Fig. 7F).

Reflecting upon the results of GAT-3 blockade upon SNr firing in isolation, an assay of the isolated effect of GAT-1 blockade upon SNr firing was deemed necessary. If blockade of GABA influx by astrocytic GAT-3 results in increased inhibition of SNr firing rate, then blockade of GABA efflux by dopaminergic GAT-1 must result in decreased inhibition of SNr firing rate (Fig. 7G, H). Indeed, the bath application of GAT-1 antagonist NNC-711 alone upon SNr neurons significantly increased mean firing rate (36% increase, p <0.01, n = 6) (Fig. 7H). Lastly, if GAT-3 transporters were mediating GABA uptake, then tonic GABA levels should be sensitive to temperature. In agreement with the hypothesis, lowering the temperature of the slice to 20-22 deg. C (from 32 deg. C) blunted the ability of gabazine to accelerate SNr spiking rate (p = 0.05, n = 7) (Fig. S4D, E).

Astrocytes have also been reported to regulate extracellular GABA concentration in other ways. In particular, astrocytes have been reported to release GABA through bestrophin1 channels (BST1) in response to an elevation in intracellular Ca^2+^ concentration (*20*). To test if SNr astrocytes utilize a similar mechanism, the SNr was injected with an AAV carrying a plasmid in which a glial fibrillary acidic protein (GFAP) promoter (*49*) fragment controlled expression of an excitatory hM3 (G_q_) DREADD construct. Previous studies have used this G_q_-linked M3 DREADD to elevate intracellular Ca^2+^ in astrocytes (*21*) (Fig. S4A). In ex vivo brain slices, bath application of the DREADD agonist CNO elevated – not suppressed – the basal spiking rate of SNr neurons recorded in cell-attached mode (p = 0.03, n = 8) (Fig. S4B). Furthermore, bath application of the BST1 channel inhibitor NPPB decreased SNr spiking rate (p = 0.03, n = 6) (Fig. S4C) – suggesting that BST1 channels in SNr astrocytes were releasing an excitatory neurotransmitter, not GABA.

## Discussion

There are three conclusions that can be drawn from the studies presented. First, activation of D2-class dopamine receptors suppressed optogenetically-induced, phasic release of GABA from GPe axon terminals synapsing on SNr neurons. Second, activation of D2-class dopamine receptors also suppressed tonic GABAergic inhibition of autonomous SNr spiking. Although a small component of this tonic GABA release was attributable to GABA release from dSPN terminals, the principal component of this tonic signaling was attributable to GAT-1-mediated dendritic release of GABA from ALDH1A1+ SNc dopaminergic neurons. Third, the dopaminergic modulation of tonic GABA signaling was attributable to D2R-mediated enhancement of GAT-3 uptake of GABA by SNr astrocytes. Taken together, these results establish that dendritic release of dopamine suppresses both phasic and tonic GABAergic inhibition of SNr spiking (Fig. 8A). Moreover, they demonstrate that the metabolism of ALDH1A1+ dopaminergic neurons, which are the most bioenergetically challenged subset of dopaminergic neurons, modulates SNr neuron spiking and, in so doing, could shape movement (Fig. 8B).

**Figure 8.**
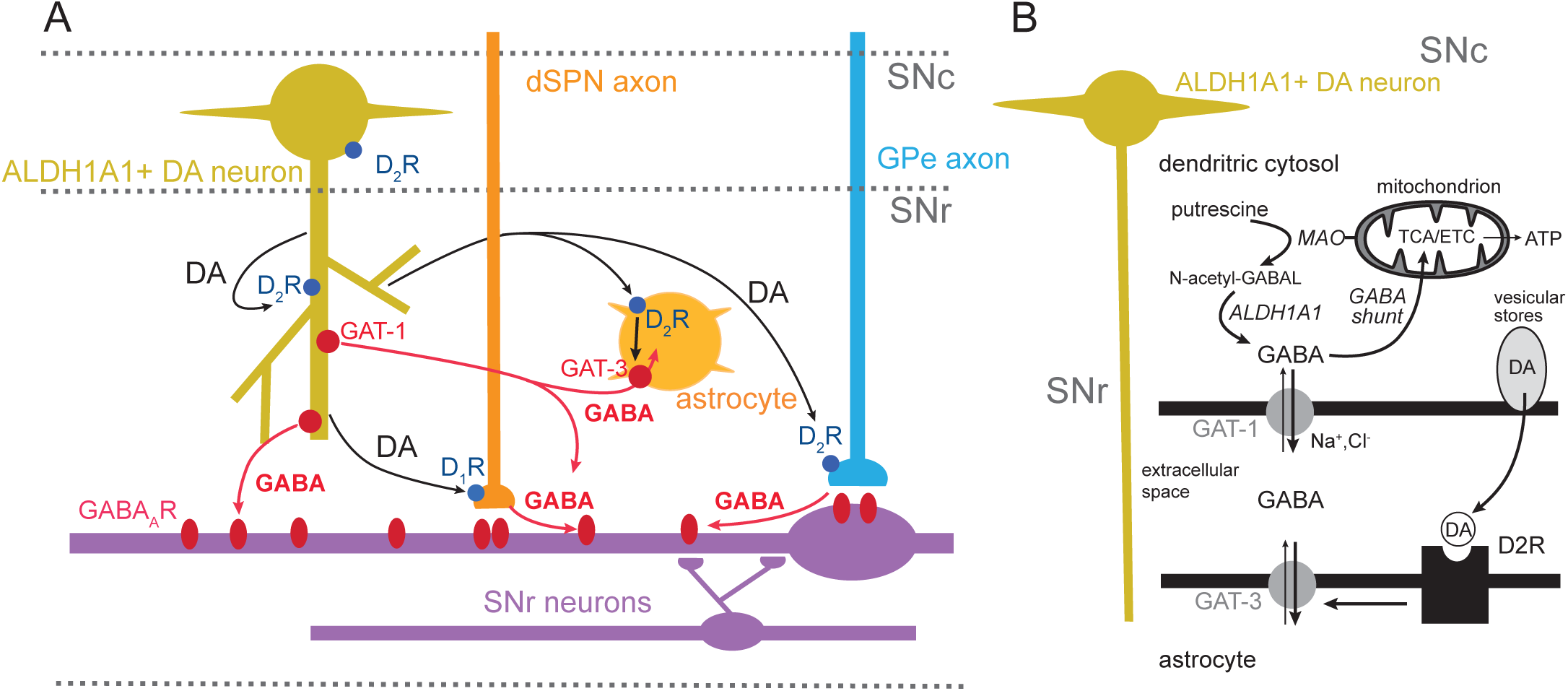
Schematic for release of DA and GABA in SNr. (A) SNc dopamine acts upon D2 receptors on GPe axons and astrocytes in the SNr. Activation of astrocytic D2Rs disinhibits SNr neurons and enhances firing rate. GPe inputs appear unrelated to tonic GABA inhibition. (B) Non-canonical synthesis of GABA feeding into the GABA shunt at SNc dopamine neuron mitochondria. Excess GABA is transported out of DA neurons into the extracellular space by GAT-1. D2R activation at astrocytes enhances GAT-3 mediated uptake of GABA from the extracellular space, indirectly disinhibiting SNr.

### Presynaptic D2-class dopamine receptors suppressed GABA release by GPe terminals

GPe GABAergic projection neurons form large perisomatic synapses on SNr neurons (*50*). Previous studies using electrical stimulation of fiber tracks, in combination with pharmacological and electrophysiological methods in ex vivo brain slices, had inferred that presynaptic D2-class dopamine receptors inhibited the release of GABA from GPe terminals on SNr neurons (*15*); this modulation was attributed to a combination of D_3_ and D_4_ dopamine receptors based upon pharmacological criteria. Given the inability of electrical stimulation to activate specific neurons and the potential heterogeneity of GABAergic inputs to SNr neurons, optogenetic approaches were used here to definitively test this inference. Although the type of D2-class receptor involved was not addressed in our studies, they do demonstrate that GPe terminals possess D2-class receptors that robustly suppress GABA release at synapses on SNr neurons.

### Tonic GABAergic signaling in SNr was largely attributable to dendritic release by SNc dopaminergic neurons

In addition to inhibiting optogenetically evoked GABA release from GPe terminals, the D2-class receptor agonist quinpirole suppressed a tonic, GABA_A_R-mediated inhibition of spontaneous spiking in SNr neurons. This tonic signaling was not diminished by chemogenetic inhibition of GPe terminals or astrocytes, pointing to an alternative origin. Our experiments identified two.

One source of tonic GABA signaling appeared to be the synaptic terminals of dSPNs, as antagonizing D1Rs, which facilitate dSPN terminal GABA release (*22–24*), modestly elevated the basal spiking rate of SNr neurons. As dendritic release of dopamine is negatively modulated by D2 autoreceptor signaling (*16, 51*), quinpirole should dis-facilitate tonic GABA release from dSPN terminals. However, the modal change in spiking rate produced by antagonizing D1Rs was small (∼10%) compared to that produced by D2R agonist quinpirole (∼50%)

Another, and more significant, source of tonic GABA signaling in SNr was ALDH1A1+ SNc dopaminergic neurons. These dopaminergic neurons reside near the border between SNc and SNr and have dendrites that course through much of the SNr. These dendrites are well-known to release dopamine; this release is activity-dependent, reliant upon vesicular mechanisms, and is negatively modulated by D2R autoreceptor signaling (*16, 51*). Our work shows that these dendrites also release GABA. In agreement with previous work examining axonal release, the dendritic generation of GABA appeared to depend upon intracellular putrescine metabolism, as it was reliant upon both MAO-B and ALDH1A1. Unlike dopamine, the transit of GABA from the dendritic cytoplasm to the extracellular space was dependent upon GAT-1 transporters, not vesicular mechanisms. Consistent with this view, GAT-1 was preferentially expressed in dopaminergic neurons expressing ALDH1A1, aligning both metabolic and release mechanisms.

What is less evident are the reasons for the synthesis and release of GABA by ALDH1A1+ dopaminergic neurons. One possibility is related to bioenergetics. ALDH1A1+ dopaminergic neurons in the ventral tier of the SNc are bioenergetically challenged because of their massive axonal arbors and sustained activity (*37–39*). Consequently, they are highly dependent upon mitochondrial ATP generation (*52, 53*). Although glycolysis-derived pyruvate is the dominant substrate for the tricarboxylic acid (TCA) cycle and the production of nicotinamide adenine dinucleotide (NADH) for the ETC (*54, 55*), intracellular GABA can provide an alternative source of reducing equivalents for the ETC when pyruvate is in short supply. Indeed, ALDH1A1+ dopaminergic neurons express both GABA transaminase and succinate dehydrogenase – the key enzymes for the GABA shunt pathway (*31, 40*). Indeed, when extracellular glucose levels were lowered, inhibition of the GABA shunt diminished mitochondrial ATP generation. Although many questions remain to be answered about this alternative metabolic pathway, its presence in dopaminergic neurons is consistent with prior work showing that neurons can engage the GABA shunt to support OXPHOS (*56*). When it is not needed, putrescine-derived GABA appears to be ’dumped’ into the extracellular space through GAT-1. Given that the reversal potential for GAT-1 GABA transport is very near the median membrane potential of dopaminergic neurons (∼-65 mV), intracellular GABA concentration would not have to rise much to drive export. In support of this metabolic hypothesis, lowering extracellular glucose pushed down tonic GABA release and the inhibition of SNr spiking.

### Coordinated modulation of SNr by GABA and DA

How do our studies inform our understanding of how SNc dopaminergic neurons regulate the activity of SNr? Although our work demonstrates that ALDH1A1+ dopaminergic neurons can release GABA from dendrites, in vivo extrasynaptic GABAergic signaling in the SNr is likely to arise from a variety of sources. As GPe neurons spike at high sustained rates, their terminals could play a prominent role in determining extrasynaptic GABA concentrations, even though this was not the case in ex vivo brain slices. Acting at D2R-class receptors, dendritically released dopamine will not only inhibit GABA release from GPe terminals, but also will enhance GAT-3 mediated astrocytic uptake of GABA ’over-flow’. Both actions should serve to accelerate SNr neuron spiking. In the classical framework, pro-kinetic dopamine release should inhibit SNr neuron spiking – not increase it. An alternative hypothesis is that the elevation in SNr neuron spiking serves to increase the information conveyed per unit time (*57*) or to increase the correlation between neurons with shared input (*58, 59*), which could be pro-kinetic. Simultaneously boosting phasic GABA release from dSPN terminals could increase the signal-to-noise ratio of movement related signals.

What do our studies tell us about the network determinants of PD symptoms? The classical model of network dysfunction argues that striatal dopamine depletion is both necessary and sufficient for the emergence of the cardinal motor symptoms of PD (*60*). But there is a growing body of evidence that extrastriatal signaling by SNc neurons plays a critical role in this transition (*61–63*). For example, in otherwise healthy rodents, disrupting dopaminergic signaling within the SNr promotes the emergence of irregular, burst mode spiking in principal neurons (*15, 64*), much like the more distributed dopamine depletion produced by 6-OHDA lesions (*65–68*). Similar alterations in the pattern of SNr spiking have been observed in primate models of PD (*69*). The emergence of the pathological bursting pattern of activity is commonly attributed to a disruption of dopaminergic modulation of afferent input to SNr neurons (*15, 70*). Our results are consistent with this hypothesis. In addition, the average spiking rate of SNr neurons drops in PD models (*59, 71*). The loss of dendritic dopamine release and impairment of GABA uptake by astrocytes will contribute to this shift, potentially compromising the ability of SNr to properly code movement.

## Materials and Methods

### Animals

All housing, breeding and procedures were performed according to the Northwestern University Animal Studies committee and in accordance with the National Institutes of Health Guide for the Care and Use of Laboratory Animals: Protocol #IS00019822 – "*Distributed circuit dysfunction underlying motor and sleep deficits in a progressive mouse model of Parkinson’s disease*", approved for up to 7988 mice. Mice were group housed with food and water provided ad libitum under a 12 h-12 h light-dark cycle and temperatures of 65-75 °F with 40-60% humidity. Male and female mice were used for experiments. No significant differences were observed between both groups. We obtained *DAT-Cre* mice (B6.SJL-Slc6a3^tm1.1(cre)Bkmn^/J) from the Jackson Laboratories. We obtained C57BL/6J and PV-Cre mice (B6.129P2-Pvalb^tm1(cre)Arbr^/J) from the Jackson Laboratories. MC1-Park mice (DAT-Cre x Ndufs2 ^-/-^ ) were bred in house (*10*). Aldh1a1-2A-iCre and Aldh1a1-2A-iCre/Dat-2A-Flpo adult mice (P>60) were generated in house.

### Stereotaxic injections

Mice were anaesthetized using an isoflurane precision vapourizer (SomnoSuite - Kent Scientific) and placed in a stereotaxic frame (David Kopf Instruments) with a Cunningham adaptor (Harvard Apparatus) to maintain anesthesia delivery throughout surgery. After exposing the skull, a small hole was bored with a micro drill bit and 200 nl of viral vector was delivered through a glass micropipette (Drummond Scientific Company) pulled on a Sutter P-97 puller. The SNr was targeted at the following coordinates (mm): anterioposterior (AP) = -3.40, mediolateral (ML) = 1.41 and dorsoventral (DV) = -4.45 (all (AP, ML, DV) relative to the bregma). For ALDH-shRNA experiments, the SNc was targeted at the following coordinates: AP = -3.0, ML = 1.2, DV = 4.2. For the DAT-FLP/ALDH1A1-Cre labeling experiments, the SNc was targeted at the following coordinates: AP = -3.16, ML = -1.5, DV = 4.00, and a volume of 300nL was injected of a mix 1:1 of AAV8 HSyn-Con/Fon-YFP (titer:2.4x10^13^GC/mL) and AAV8 Ef1a-Coff/Fon-mCherry (titer: 2.2X10^13^GC/mL). The GP was targeted at the following coordinates: AP = -0.34, ML = 1.81, DV = 3.61, and AP = -0.82, ML = 2.33, DV = 3.72.

Source of coordinates and diagrams: Allen Mouse Brain Atlas

### Recombinant AAV vectors

The following AAV viruses were used in this study: AAV9-hSyn-hM4Di-mCherry (Addgene; RRID:Addgene_50475), AAV9-hSyn-DIO-hM4Di-mCherry (Addgene; RRID:Addgene_44362), AAV2/5-GfAABC_1_D-hM3Dg-mCherry (Addgene; RRID:Addgene_50478), AAV-CMV-Aldh1a1-shRNA (Stanford, (*25*)), AAV8 HSyn-Con/Fon-YFP (addgene: #55650-AAV8, titer:2.4x10^13^GC/mL) and AAV8 Ef1a-Coff/Fon-mCherry (addgene: #137134-AAV8, titer: 2.2X10^13^GC/mL) , AAV9-TH-GW1-Perceval-SV40pA (addgene: #246242), and AAV9-EF1a-DIO-GW1-PercevalHR-WPRE (addgene: #248353).

### Ex vivo brain slice preparation

Mice were anaesthetized with a mixture of ketamine (50 mg kg^-1^) and xylazine (4.5 mg kg-1) then transcardially perfused with ice-cold, oxygenated modified artificial cerebrospinal fluid (aCSF) containing 210 mM sucrose, 2.5 mM KCl, 1.25 mM NaH_2_PO_4_, 25 mM NaHCO_3_, 0.5 mM CaCl_2_, 10 mM MgSO_4_ and 10 mM glucose, pH 7.3 (315–320 mOsm l^−1)^. Once perfused, the brain was rapidly removed and either sagittal or coronal slices containing the SNr (thickness, 240 µm) were sectioned using a vibratome (VT1200S Leica Microsystems). Brain slices were incubated in oxygenated modified aCSF containing 124 mM NaCl, 4.5 mM KCl, 25 mM NaHCO_3_, 1.25 mM NaH_2_PO_4_, 1.2 mM CaCl_2_, 1.8 mM MgCl_2_ and 10 mM glucose at 34 °C for 30 min, then at room temperature for another 30 min before the experiments. All solutions were pH 7.4, 310-320 mOsm l^-1^ and were continually bubbled with 95%O_2_/5% CO_2_. Experiments were performed at 32-33 °C.

### Fast-scan cyclic voltammetry (FSCV)

Extracellular dopamine release was monitored by fast-scan cyclic voltammetry recordings performed in 300 μm coronal sections of dorsal striatum using carbon-fiber microelectrodes (7 μm diameter carbon fiber extending 50∼100 μm beyond the tapered glass seal). Cyclic voltammograms were measured with a triangular potential waveform (-0.4 to +1.3 V vs Ag/AgCl reference electrode, 400 V/s scan rate, 8.5 ms waveform width) applied at 100 ms intervals. The carbon fiber microelectrode was held at -0.4 V between scans. Cyclic voltammograms were background-subtracted by averaging 10 background scans. Each striatal hemisphere was recorded in both dorsal lateral and dorsal medial striatum, and the order of recording was counterbalanced across slices. Dopaminergic axon terminals were stimulated locally (100-200μm from carbon fiber) with an electrode at 100 μA. First, each recording site received 2 electrical stimulations separated by 15seconds, after 2 minutes of recovery time, the same site received a train of stimulation (5 pulses at 25Hz). Evoked dopamine release and subsequent oxidation current was detected and monitored using the TarHeel CV system. Changes in dopamine concentration by optical stimulation were quantified by plotting the peak oxidation current of the voltammogram over time. Carbon fiber microelectrode was calibrated at the end of each day of experiments to convert oxidation current to dopamine concentration using 10 μM dopamine in ACSF.

### 2PLSM optical workstation and PercevalHR experiments

The laser scanning optical workstation embodies an *Ultima* dual-excitation-channel scan head (Bruker Nano Fluorescence Microscopy Unit). The foundation of the system is the Olympus BX-51WIF upright microscope with a LUMPFL 60X/1.0NA water-dipping objective lens. The automation of the XY stage motion, lens focus, and manipulator XYZ movement was provided by FM-380 shifting stage, axial focus module for Olympus scopes, and manipulators (Luigs & Neumann). Cell visualization and patching were made possible by a variable magnification changer, calibrated to 2x (100 µm FOV) as defined by the LSM bright-field transmission image, supporting a 1 Mpixel USB3.0 CMOS camera (DCC3240M; Thor Labs) with ∼30% quantum efficiency around 770nm.

Olympus NIR-1 bandpass filter, 770nm/100nm, and *μManager* (*72*) software were used with the patch camera. The electrical signals were sent and collected with a 700B patch clamp amplifier and *MultiClamp Commander* software with computer input and output signals were controlled by *Prairie View* 5.3-5.5 (RRID:SCR_017142) using a National Instruments PCI6713 output card and PCI6052e input card.

The 2P excitation (2PE) imaging source was a Chameleon Ultra1 series tunable wavelength (690-1040 mm, 80 MHz, ∼250 fs at sample) Ti: sapphire laser system (Coherent Laser Group); the excitation wavelength was selected based on the probe being imaged (see below). Each imaging laser output is shared (equal power to both sides) between two optical workstations on a single anti-vibration table (TMC). Workstation laser power attenuation was achieved with two Pockels’ cell electro-optic modulators (models M350-80-02-BK and M350-50-02-BK, Con Optics) controlled by *Prairie View* 5.3–5.5 software. The two modulators were aligned in series to provide enhanced modulation range for fine control of the excitation dose (0.1% steps over five decades), to limit the sample maximum power, and to serve as a rapid shutter during line scan or time series acquisitions.

The 2PE generated fluorescence emission was collected by non–de-scanned photomultiplier tubes (PMTs). Green channel (490–560 nm) signals were detected by a Hamamatsu H7422P-40 select GaAsP PMT. Red channel (580–630 nm) signals were detected by a Hamamatsu R3982 side on PMT. Dodt-tube-based transmission detector with Hamamatsu R3982 side on PMT (Bruker Nano Fluorescence) allowed cell visualization during laser scanning. Scanning signals were sent and received by the NI PCI-6110 analog-to-digital converter card in the system computer (Bruker Nano Fluorescence). All XY images were collected with pixel size of 0.195 µm and pixel dwell time of 12 µs and a frame rate of 3 to 4 fps.

For the excitation ratio probe PercevalHR, ATP/ADP ratio measurements of two excitation wavelengths, 950 nm for ATP and 820 nm for ADP, were used in rapid succession for each acquisition time point. Two time series of 5 frames (1.25-1.7 s long) were acquired for each wavelength. After observing a stable baseline ratio for 4 acquisitions, pharmacological manipulations were used to measure the contribution of different pathways to cellular bioenergetics. Time series were analyzed offline with FIJI (RRID: SCR_002285) (*73*), a cytosolic and a background ROIs were measured, the background signal was subtracted, and the F950/F820 ratio was calculated for each ratio pair time point.

For PercevalHR experiments, modified protective aCSFs were used for perfusion and slicing, with similar results: sucrose/low Ca^2+^ slicing solution contained: 49.14 mM NaCl, 2.5 mM KCl, 1.43 mM NaH2PO4, 25 mM NaHCO3, 25 mM glucose, 99.32 mM sucrose, 10 mM MgCl2 and 0.5 mM CaCl2; glycerol slicing solution contained: 235 mM glycerol, 2.5mM KCl, 1.25mM NaH_2_PO_4_, 25 mM NaHCO_3_, 2 mM CaCl_2_, 1 mM MgCl_2_, 10 mM glucose. Experiments were performed using 1.5 mM glucose ACSF, containing 136.75 mM NaCl, 2.5mM KCl, 1.25mM NaH_2_PO_4_, 25 mM NaHCO_3_, 2 mM CaCl_2_, 1 mM MgCl_2_, 1.5 mM glucose, in order to maximize OXPHOS. 2-DG ACSF contained 135.75 mM NaCl, 2.5mM KCl, 1.25mM NaH_2_PO_4_, 25 mM NaHCO_3_, 2 mM CaCl_2_, 1 mM MgCl_2_, 3.5 mM 2-deoxyglucose (Sigma-Aldrich). All solutions were pH 7.4 and ∼310 mOsm, and continually bubbled with 95% O_2_ and 5% CO_2_.

### Analysis and statistics

Electrophysiology data were generally analyzed using Clampfit 10.3 software (Molecular Devices; https://www.moleculardevices.com/products/axon-patch-clamp-system/acquisition-and-analysis-software/pclamp-software-suite. Data are summarized using box plots showing median values, first and third quartiles, and range, unless otherwise specified. Statistical analysis was performed with Mathematica 13 (Wolfram; https://www.wolfram.com/mathematica/?source=nav) and GraphPad Prism 8 (GraphPad Software; https://www.graphpad.com/). Probability threshold for statistical significance was *P* < 0.05.

## Acknowledgments

We thank Dr. Bal Khakh from the UCLA David Geffen School of Medicine for his insights on the astrocyte experiments.

## Author contributions

Conceptualization: DVS, CW, DJS

Methodology: DVS, DJS

Investigation: DVS, OAM, DDAR, KK, AP, NV, ZX, TT, EZ, AISDS

Visualization: DVS, OAM, DDAR, KK, NV, ZX, EZ, AISDS

Supervision: JD, RA, DJS

Writing—original draft: DVS, DJS

Writing—review & editing: DVS, OAM, DDAR, KK, AISDS, EZ, DJS,

## Data, Code, & Materials Availability Statement

All datasets, protocols, key lab materials used, and code generated in this study are listed in a key resource table (10.5281/zenodo.17296457) alongside their public persistent identifiers. This study did not generate new materials.

