## Supplemental Figure Legends for "Astrocytes mediate the dopaminergic modulation of tonic GABAergic signaling in substantia nigra"

**Figure. S1. Delta in firing rate after gabazine weakly correlated with base rate.**

Summary data and linear regression of a weak positive correlation between basal discharge rate and the effect of gabazine R2 = 0.11, n = 39

**Figure S2. D1R antagonist significantly elevates SNr spike rate.** (A) Example trace and summary data of cell attached spiking from SNr neurons before and after bath application of SCH39166. Paired T-test, n = 12, p = 0.02. (B) Bath applied D2R agonist elevates SNr firing rate significantly more than bath applied D1R antagonist. Unpaired T-test, n = 8 D2R / 12 D1R, p = 0.03.

**Figure S3. GiDREADD Inhibits SNc Neurons.** Example trace and summary data: CNO activation of Gi-DREADD inhibits the firing rate of SNc neurons (mean reduction of 19%, p = 0.02, n = 6).

**Figure S4. Astrocyte modulation of SNr spiking is unrelated to Tonic GABA**
(A) Proposed schematic of Gq-DREADD expression in Astrocytes. CNO application triggers intracellular calcium release and modulates conductance of bestrophin 1 channels. (B) Example trace and summary data of cell attached spiking from SNr neurons before and after bath application of DREADD agonist CNO. Paired T-test, n = 8, p = 0.03. (C) Example trace and summary data of cell attached spiking from SNr neurons before and after bath application of BST1 antagonist NPPB. Paired T-test, n = 6, p = 0.03. (D) Example trace and summary data of SNr neurons examined in cell-attached mode at room temperature (20-22 o C) exhibiting no sensitivity to gabazine (p = 0.05, n = 7). (E) Summary data comparison of the SNr neuron firing rate fold change following gabazine treatment at physiological (30oC) and room temperature (20oC) Mann- Whitney test, n = 44/7, p < 0.0001.

**Figure S5: Table of GAT (1-4) expression in SNC dopaminergic cells.**

Gonzalez-Rodriquez et.al. 2021 DOI: 10.1038/s41586-021-04059-0

**Figure S6. Deletion of ALDH1A1 does not affect basal dopamine release.** (A) Schematic of the FSCV experiment in coronal striatal slices. Inset: 2D voltammogram showing oxidation and reduction peaks of DA. (B) Representative 3D voltammograms from paired pulse stimulation; x axis: recording time; y axis: applied potential; color map: recorded current. (C-D) Quantification of peak DA release evoked by 1-pulse (C) and 5-pulse (D) stimulation. 1 pulse data are from the first pulse of the paired pulse represented by (B). (E) Quantification of the 15s ISI paired pulse ratio represented by (B).

**Figure S7. RNA Scope in situ hybridization assay of D2R colocalization with S100 beta within SNr Astrocytes.** Overall, 35 +/- 15% (Mean +/- SD) astrocytes are D2R-positive (median 33%); n = 3 mice/516 astrocytes. (A) Low power coronal image of midbrain with D2R and S100 beta labeling. (B,C) High power inset images of SNr with DAPI stain (B) and without DAPI stain (C)

**Figure S8. Low external glucose blocks tonic GABA.** (A) Summary data: When extracellular glucose concentration is lowered to 2.5 mM, gabazine does not significantly increase SNr spiking. Paired T-test, n = 8, p = 0.10 . (B) Summary data from Figure 3B included for comparison.

**Figure S9. GABA is used as a bioenergetic substrate for mitochondrial OXPHOS
in dopaminergic neurons.** (A) Representative 2-PLSM image (average of 5 frames) of a
SNc dopaminergic neuron expressing PercevalHR; scale bar, 10 μm. (B) Representative time-lapse trace of a PercevalHR experiment. The ratio between the fluorescence intensity
when exciting the sample with 950 nm and 820 nm wavelengths is measured at intervals of
5 minutes, and the sequential pharmacological manipulations are used to dissect the
contribution of different pathways to cellular bioenergetics. Bath application of vigabatrin (
20 μM) and oligomycin (10 μM) inhibits GABA transaminase (GABA-T) and mitochondrial Complex V, respectively. Glucose replacement with 2-deoxyglucose (3.5mM) blocks glycolysis. (C) Box plots summarizing the effect of GABA-T inhibition on SNc neurons bioenergetics. Wilcoxon matched pairs signed rank test, p = 0.002. Experiments were performed on wild-type mice using a probe under a tyrosine hydroxylase (TH) promoter (open symbols, n = 7, N = 7)
or on Aldh1a1-cre mice using a probe under a cre-dependent promoter (full symbols, n = 3, N = 3) for higher sub-regional selectivity, and subsequently pooled together. (D) Box plots comparing the 2-DG sensitive (glycolytic) contribution to cellular bioenergetics in controls and vigabatrin-treated cells; n = 6, N = 5; n = 10, N = 10 respectively; Mann-Whitney test, p = 0.367. (E) Box-plots showing that vigabatrin decreases the oligomycin-sensitive (OXPHOS) contribution to cellular bioenergetics; n = 6, N = 5 for controls, n = 10, N = 10 for vigabatrin-treated cells. Mann-Whitney test, p = 0.0017. Box-plots represent median (thick line), interquartile range, and minimum and maximum (whiskers). Individual points are also indicated.
