## Supplementary figures and images for "Astrocytes mediate the dopaminergic modulation of tonic GABAergic signaling in substantia nigra"

### Figure S1

**Figure S1. Delta in firing rate after gabazine weakly correlated with Base rate**

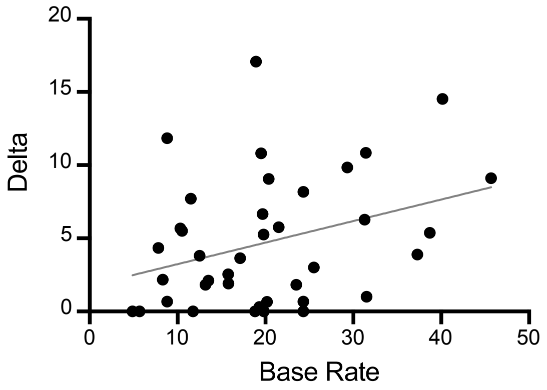

### Figure S2

**Figure S2. D1R antagonist significantly elevates SNr spike rate.**

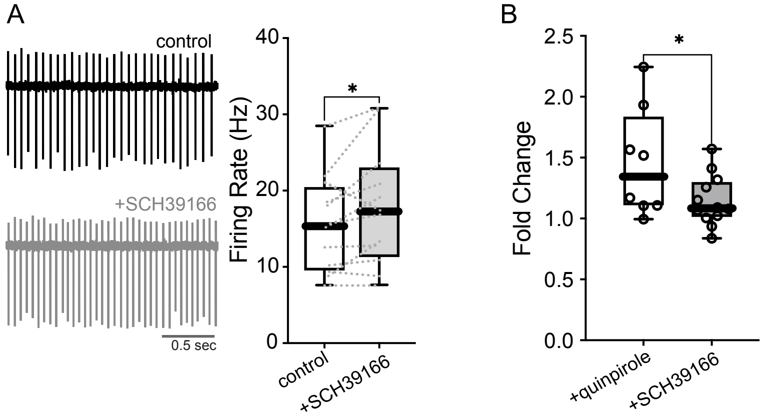

### Figure S3

**Figure S3. GiDREADD Inhibits SNc Neurons**

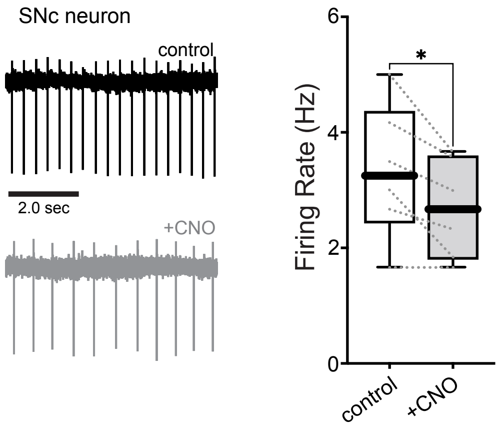

### Figure S4

**Figure S4. Astrocyte modulation of SNr spiking and Tonic GABA**

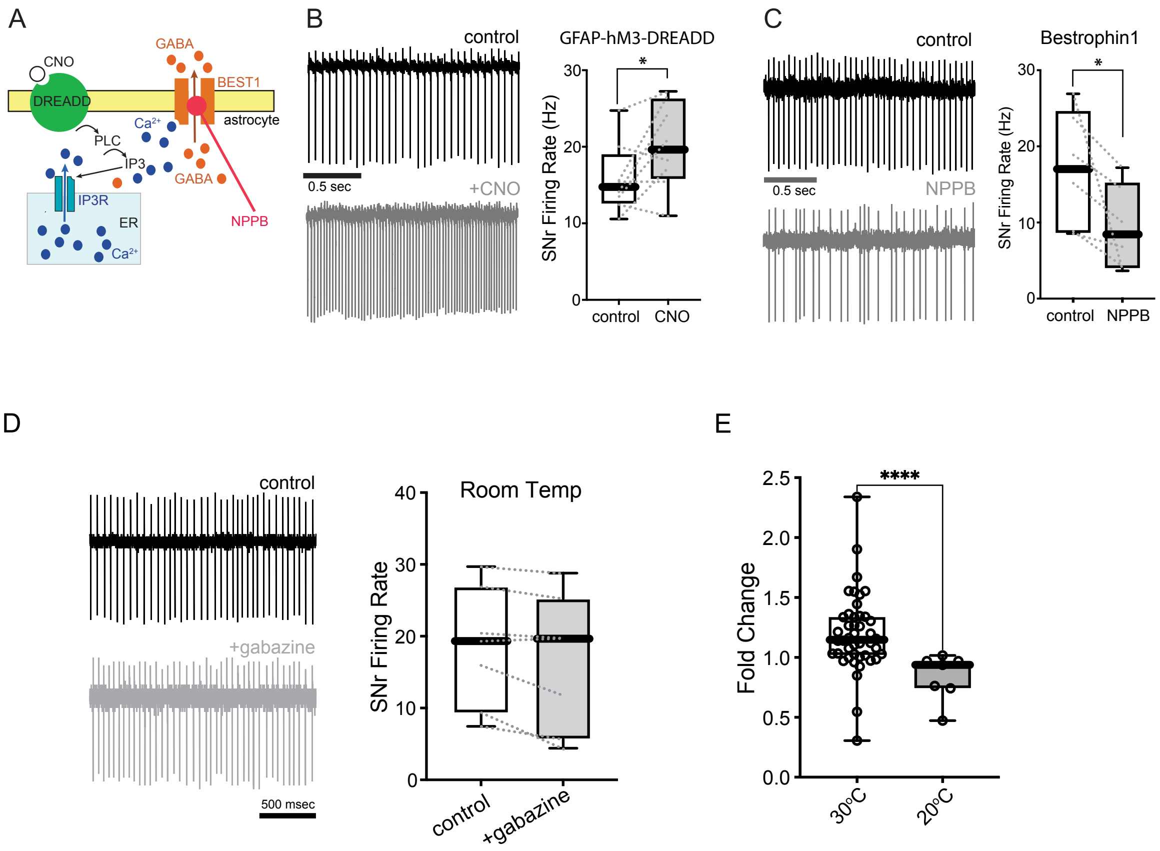

### Figure S6

**Figure S6. Deletion of ALDH1A1 does not impact dopamine release.**

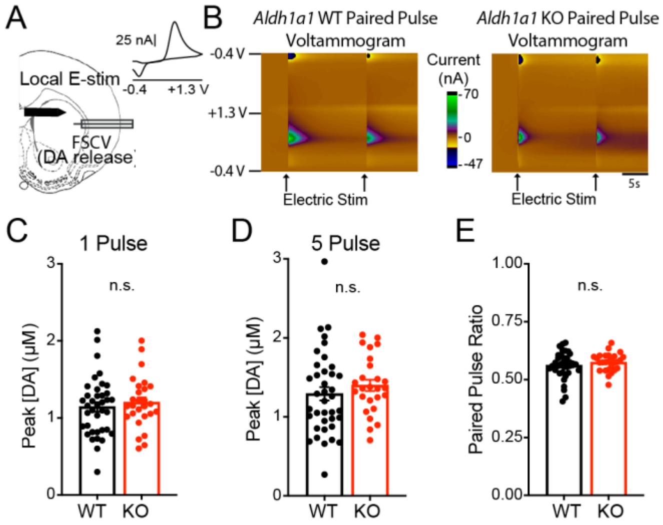

### Figure S7

**Figure S7. Quantification of D2R expression in SNr Astrocytes**

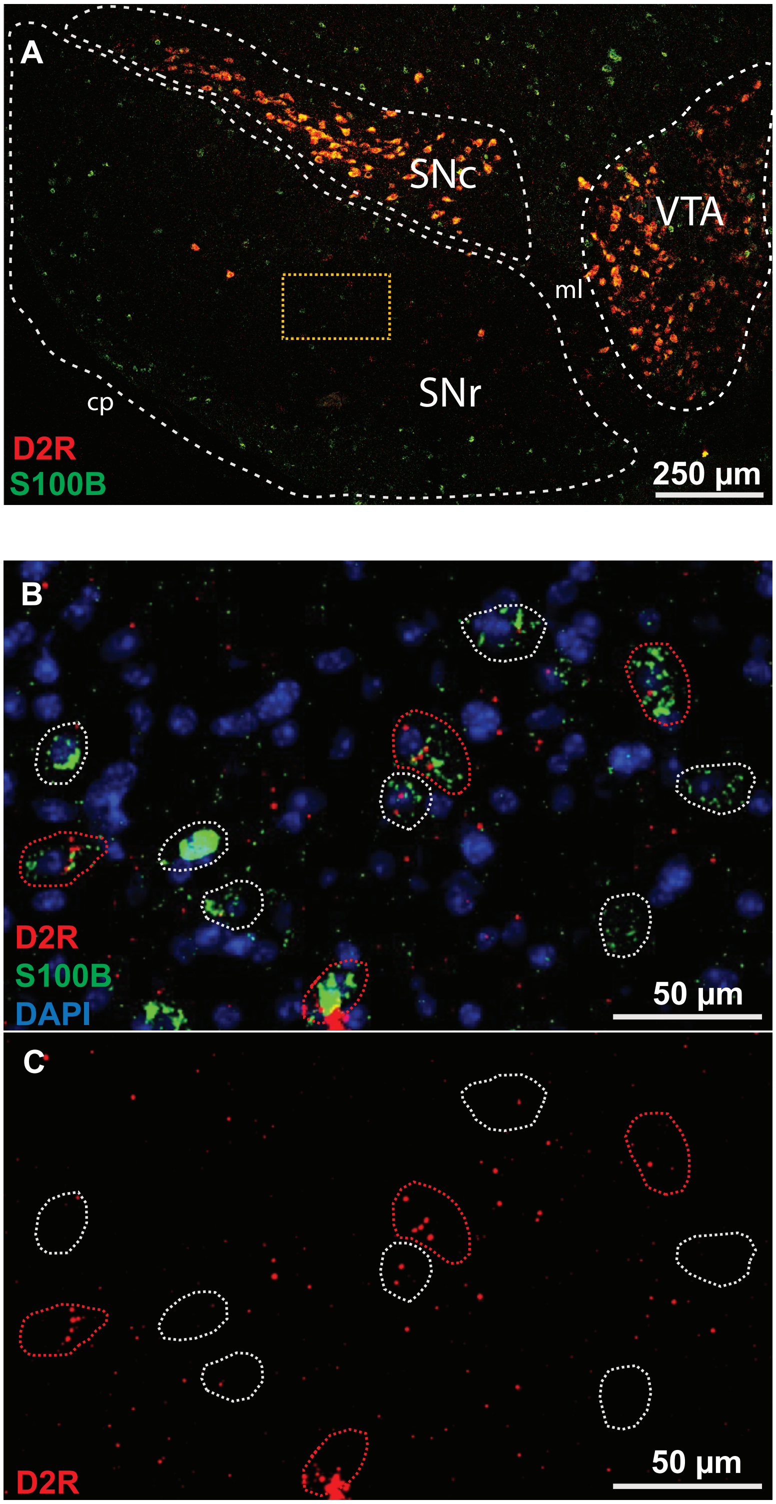

### Figure S8

**Figures S8. Gabazine does not facilitate SNr firing rate in 2.5 mM Glucose**

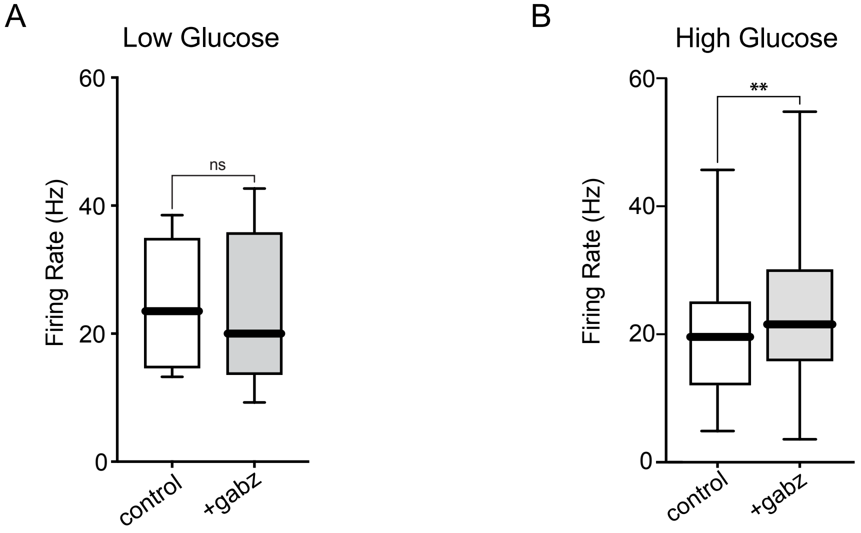

### Figure S9

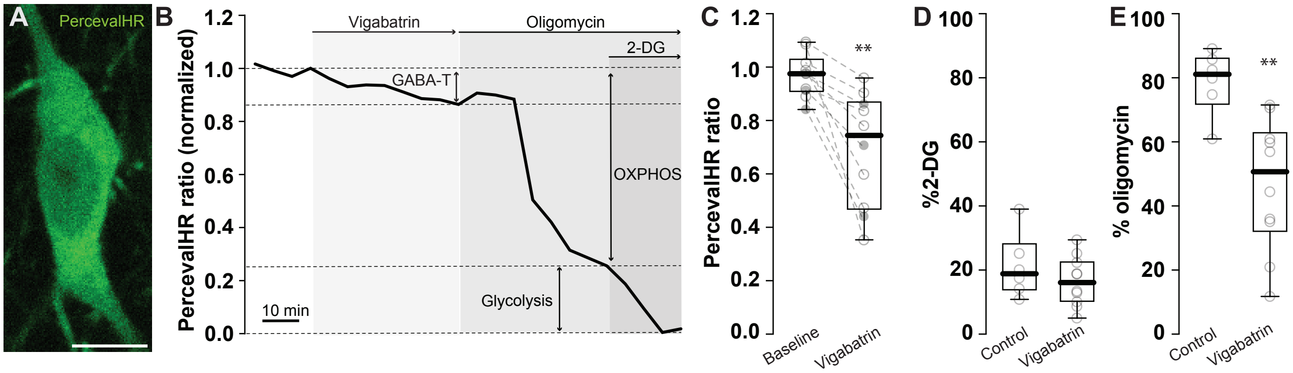
