## Supplementary material for "Astrocytes mediate the dopaminergic modulation of tonic GABAergic signaling in substantia nigra": Figure S5

**Figure S5. Table of RNAseq reads of solute carrier family 6 genes from SNc Dopamine neurons**

| Gene Symbol | Description | Reads |
|---|---|---|
| Slc6a1 | solute carrier family 6 (neurotransmitter transporter, GABA), member 1 | 2118.494118 |
| Slc6a12 | solute carrier family 6 (neurotransmitter transporter, betaine/GABA), member 12 | 0 |
| Slc6a11 | solute carrier family 6 (neurotransmitter transporter, GABA), member 11 | 1444.866233 |
| Slc6a13 | solute carrier family 6 (neurotransmitter transporter, GABA), member 13 | 13.86342026 |
